# Bone-Specific Enhancement of Antibody Therapy for Breast Cancer Metastasis to Bone

**DOI:** 10.1101/2021.08.31.457412

**Authors:** Zeru Tian, Chenfei Yu, Weijie Zhang, Kuan-lin Wu, Ruchi Gupta, Zhan Xu, Ling Wu, Yuda Chen, Xiang H. -F. Zhang, Han Xiao

## Abstract

Therapeutic antibodies have gone a long way toward realizing their clinical potential and have become very useful for treating a variety of pathologies. Despite the rapid evolution of therapeutic antibodies, their clinical efficacy in treatment of bone tumors has been hampered by the inadequate pharmacokinetics and poor bone tissue accessibility of these large macromolecules. Here, we show that engineering therapeutic antibodies to include bone-homing peptide sequences dramatically enhances their concentration in the bone metastatic niche, resulting in significantly reduced survival and progression of breast cancer bone metastases. To enhance the bone tumor-targeting ability of engineered antibodies, we introduced varying numbers of a bone-homing peptide into permissive internal sites of the anti-HER2 antibody trastuzumab. Compared to the unmodified antibody, the engineered bone-targeting antibodies have similar pharmacokinetics and *in vitro* cytotoxic activity against HER2-positive cancer cells, but exhibit improved bone tumor distribution *in vivo*. Accordingly, in xenograft models of breast cancer metastasis to bone sites, engineered antibodies with enhanced bone specificity exhibit increased inhibition of both initial bone metastases and secondary multi-organ metastases from bone lesions. Furthermore, this engineering strategy is also applied to prepare bone-targeting antibody-drug conjugates with enhanced therapeutic efficacy. These results demonstrate that adding bone-specific targeting to antibody therapy results in robust delivery of therapeutic antibodies to the bone tumor niche. This provides a powerful strategy for overcoming inadequate treatment of bone cancer and the development of potentially acquired resistance to therapy.

## Introduction

Antibody-based therapies entered the clinic over 30 years ago and have become the mainstream therapeutic option for patients with malignancies,^1,2^ infectious diseases,^3,4^ and transplant rejection.^5^ Compared with traditional chemotherapy, these biotherapeutics preferentially target cells presenting tumor-associated antigens, resulting in improved treatment outcomes and reduced side effects.^6,7,8,9^ Despite their high affinity for tumor antigens, poor tumor tissue penetration and heterogeneous distribution of therapeutic antibodies in brain and bone have significantly limited their efficacy in treating diseases in these tissues. Failure to deliver efficacious antibody doses throughout the tumor in these tissues leads not only to treatment failure, but also to development of acquired drug resistance.^10^ Exposure to subtherapeutic antibody levels has been shown to facilitate tumor cell ability to evade antibody-mediated killing.^11,12^ Furthermore, attempts to ensure effective concentrations of antibodies in the tumor niche usually leads to high concentrations in other tissues, resulting in adverse systemic side effects that may limit or exclude use of the therapeutic. Thus, strategies to improve tumor penetration and distribution of antibodies in a specific tissue following systemic delivery are crucial for optimizing the clinical potential of these agents.

Despite a 5-year survival rate greater than 90%, between 20-40% of breast cancer survivors will eventually experience metastases to distant organs, even years after the initial treatment.^13^ Bone is the most frequent tissue for breast cancer metastases.^14,15^ Dosing the bone microenvironment has proved to be difficult due to the relatively low density of vascularization and the presence of physical barriers to penetration. Antibody-based therapies face special distribution difficulties due to the large molecular size of these agents. Thus, therapeutic antibodies that exhibit excellent efficacy for the treatment of primary mammary tumors yield only suboptimal responses in patients with bone metastases. For example, the trastuzumab (Herceptin) antibody that successfully targets human epidermal growth factor receptor 2 (HER2) in primary breast tumors has been evaluated as a treatment option for patients with metastatic breast cancer. Although some breast cancer patients benefit from these treatments, a large number of breast cancer patients with bone metastasis experience further tumor progression within one year, and few patients achieve prolonged remission.^16^ Thus, the efficacy of therapeutic antibodies appears to be particularly limited in the case of bone metastases.

Bones are composed primarily of hydroxyapatite (HA) crystals, the insoluble salts of calcium and phosphorus. The restricted distribution of HA in hard tissues such as bone makes it an attractive target for selective bone targeting. Nature has evolved a variety of HA-binding proteins, including sialoprotein and osteopontin, that provide sites for cell anchorage and for modulating the bone mineralization process. Interestingly, sequence analysis reveals that these proteins have repeating sequences of acidic amino acids that represent possible bone binding sites (**Fig. 1A**).^17^ Short bone-homing peptides consisting of aspartic acid (Asp) have been tested for specific delivery of small molecules, microRNAs, and nanoparticles to the bone niche.^18–22^ These short peptides have been shown to favor binding to HA surface with higher levels of crystallinity. This surface is characterized by the presence of bone resorption surfaces and are known as the osteolytic bone metastatic niche.^23^ Formation of osteolytic bone lesions is driven by paracrine crosstalk among cancer cells, osteoblasts, and osteoclasts.^22–26^ Specifically, cancer cells secrete molecules such as parathyroid hormone-related protein (PTHrP) and interleukin 8 that stimulate osteoclast formation directly or indirectly by acting to modulate the expression of osteoblast genes such as receptor activator of nuclear factor-*κ*B ligand (RANKL) and osteoprotegerin (OPG). The consequent increase in bone resorption leads to release of growth factors (e.g., IGF1) that reciprocally stimulate tumor growth. Thus, selective delivery of therapeutic agents to the bone metastatic niche has the potential to interrupt this vicious osteolytic cycle. Here, we report a general strategy for engineering antibodies with bone-homing peptides capable of enhanced targeting of bone tumors. Following the insertion of bone-homing peptide sequences into anti-tumor antibodies, we demonstrate the engineered antibodies and antibody-drug conjugates with a moderate bond-targeting capability exhibit optimal efficacy to inhibit breast cancer metastases as well as multi-organ secondary metastases in xenograft models (**Fig. 1B**).

**Figure 1.**
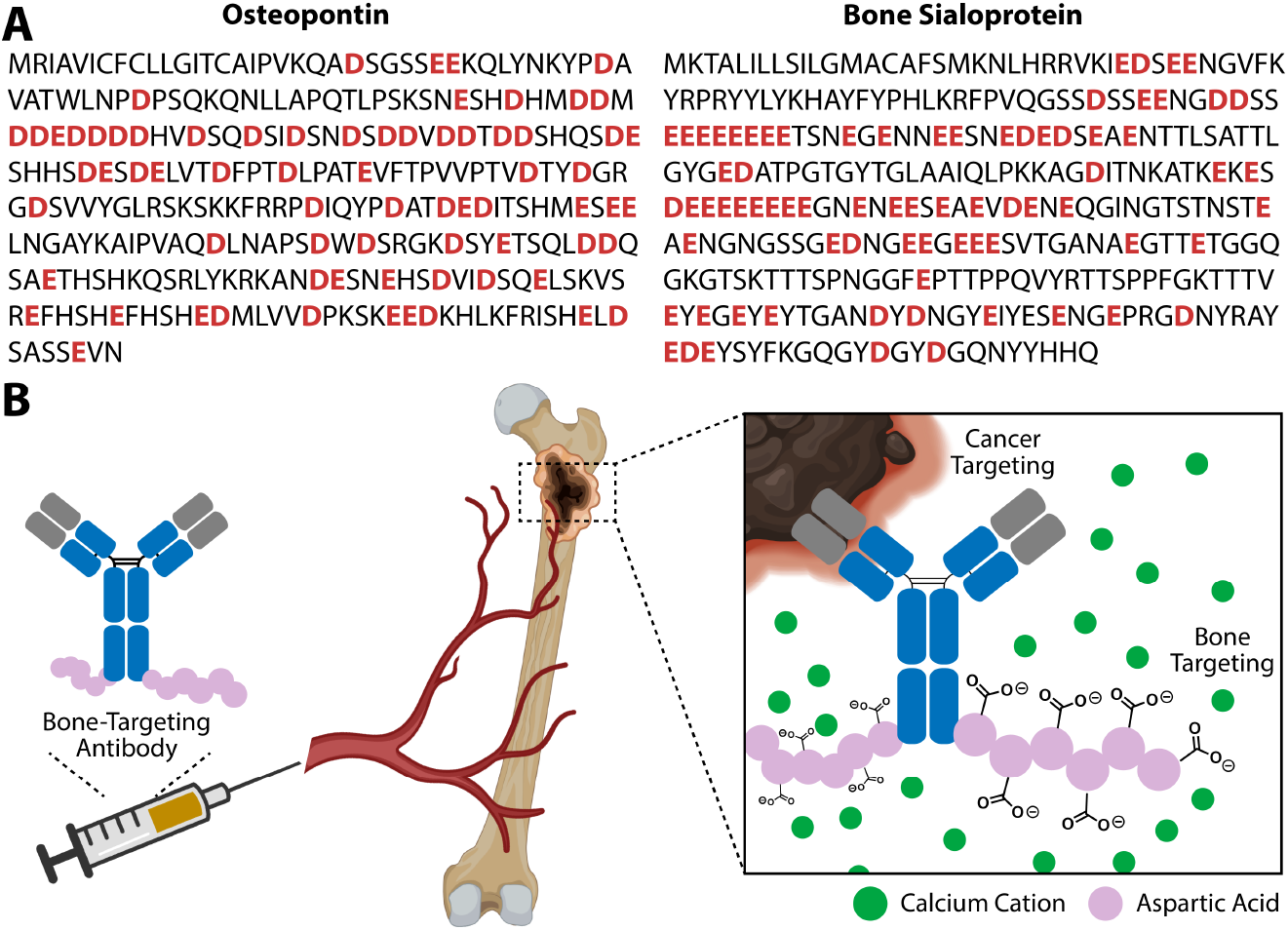
**(A)** Protein sequences of hydroxyapatite-binding proteins. **(B)**Therapeutic antibodies can be specifically delivered to the bone by introducing bone-homing peptide sequence that bind to the bone hydroxyapatite matrix.

## Results

### Modifying Trastuzumab with Bone-Homing Peptides

To harness the power of bone-homing peptide for selective delivery of antibodies to bone cancer sites, we first engineered a library of antibodies carrying the bone-homing peptide *L*-Asp_6_ at various sites within the immunoglobulin molecules. Our design principle was to install the bone-homing sequence at sites that would be minimally disruptive to the native IgG structure and function, yet would allow retention of the peptide’s high affinity for bone matrix. Based on the crystal structure of an IgG1 monoclonal antibody, we inserted the *L*-Asp_6_ peptide into permissive internal sites in the trastuzumab light chain (LC, A153), heavy chain (CH1, A165), and C-terminus (CT, G449) to yield Tras-LC, Tras-CH1, and Tras-CT, respectively (**Fig. 2A**). These internal sites have been shown to be stable to peptide insertion by screening an antibody peptide-placement library.^27^ To modulate the bone tumor-targeting ability of engineered antibodies, we also varied the number of bone-homing peptide sequences per immunoglobulin molecule, generating trastuzumab species with two (Tras-LC/CT, Tras-CH1/CT, and Tras-LC/CH1) and three *L*-Asp_6_ peptide sequences (Tras-LC/CH1/CT). The resulting seven constructs were expressed in ExpiCHO-S cells by transient transfection, followed by purification of immunoglobulins using protein G chromatography and analysis of expressed proteins by SDS-PAGE. To our delight, all these antibody mutants were expressed in good yield (50 - 100 mg/L). SDS-PAGE and ESI-MS analysis confirmed the successful insertion of the bone-homing peptides (**Fig. 2B, 2C, S1-8**). Among the antibody variants, the Tras-LC/CH1 species containing *L*-Asp_6_ peptide sequences in both the light chain and heavy chain exhibited significant aggregation. Thus, this antibody mutant was not be studied further.

**Figure 2.**
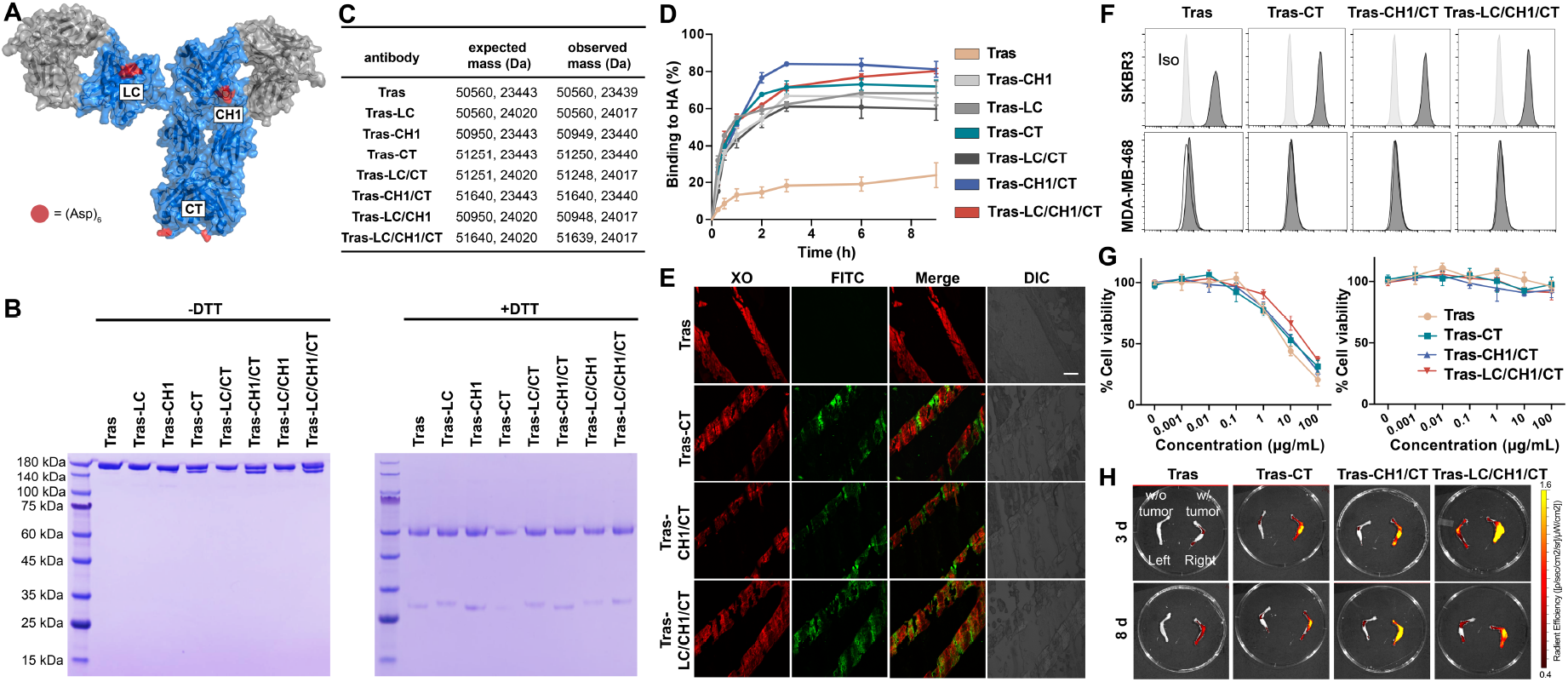
Preparation and characterization of bone-targeting antibodies. **(A)** We inserted the bone-homing peptide at three locations: light chain (LC), heavy chain (CH1), and c-terminus (CT). **(B)** SDS-PAGE analysis of bone-targeting antibodies in the absence (left) and presence (right) of the reducing reagents. **(C)** Mass spectrometry analysis of bone-targeting antibodies. **(D)** Binding kinetics of Tras, Tras-CH1, Tras-LC, Tras-CT, Tras-LC/CT, Tras-CH1/CT, and Tras-LC/CH1/CT to hydroxyapatite (HA). **(E)** Differential bone targeting ability of Tras and bone-targeting antibodies. Nondecalcified bone sections from C57/BL6 mice were incubated with 50 µg/mL Tras or bone-targeting antibodies overnight, followed by staining with fluorescein isothiocyanate (FITC)-labeled anti-human IgG and 4 µg/mL xylenol orange (XO, known to label bone), Scale bars, 200 µm. **(F)** Flow cytometric profiles of Tras, Tras-CT, Tras-CH1/CT, and Tras-LC/CH1/CT binding to SK-BR-3 (HER2+++) and MDA-MB-468 (HER2-) cells. **(G)** *In vitro* cytotoxicity of Tras, Tras-CT, Tras-CH1/CT, and Tras-LC/CH1/CT against SK-BR-3 and MDA-MB-468 cells. **(H)** *Ex vivo* fluorescence images of lower limbs of athymic nude mice bearing MDA-MB-361 tumors 72 h or 120 h after the retro-orbital injection of Cy7.5-labeled Tras, Tras-CT, Tras-CH1/CT, and Tras-LC/CH1/CT antibodies. Tumor cells were inoculated into the right tibiae of nude mice via para-tibial injection.

### *In vitro* Evaluation of Bone-Targeting Antibodies

With the bone-targeting antibody variants in hand, we initially used a HA binding assay to examine their binding to mineralized bone. Briefly, bone-targeting antibody species were incubated with HA for varying lengths of time, and unbound antibody remaining in solution was measured using a UV-Vis spectrophotometer. As shown in **Fig. 2D**, unmodified Tras exhibited only slight affinity for HA, while *L*-Asp_6_ peptide-modified antibodies bound to HA in a time-dependent manner. As expected, antibodies with multiple *L*-Asp_6_ peptides, namely Tras-CH1/CT and Tras-LC/CH1/CT, exhibit the highest HA binding capacity, with over 80% of the antibody bound after 9 h of incubation (**Table S1**). Regarding the three antibody species containing single *L*-Asp_6_ peptides, the C-terminal construct (Tras-CT) exhibits the highest HA binding capacity. Accordingly, fluorescein isothiocyanate (FITC)-labeled Tras, Tras-CT, Tras-CH1/CT, and Tras-LC/CH1/CT were used to stain non-decalcified bone sections from C57BL/6 mice. Sections treated with unmodified Tras exhibited no fluorescence after overnight incubation **(Fig. 2E)**. In contrast, we observed FITC signals in all the sections stained with the three *L*-Asp_6_ peptide-containing variants. These FITC antibody signals correlated well with the xylenol orange (XO) signal from the bone **(Fig. 2E, S9)**.

To demonstrate that insertion of the *L*-Asp_6_ sequence has negligible influence on Tras antibody binding and specificity, FITC-labeled Tras, Tras-CT, Tras-CH1/CT, and Tras-LC/CH1/CT antibodies were tested for binding to HER2-positive and negative cell lines. Flow cytometry reveals that, while none of these antibodies bind to HER2-negative MDA-MB-468 cells, each of the bone-targeting antibodies bind to HER2-expressing SK-BR-3 cells with a *K*_*d*_ similar to that of unmodified Tras (7.09 nM) (**Fig. 2F, S10-19, and Table S2)**. The Tras-CT, Tras-LC, Tras-CH1, Tras-CH1/CT, Tras-CH1/CT and Tras-LC/CH1/CT species have slightly higher K_d_ values than Tras (14.60 nM, 19.39 nM, 12.99 nM, 19.47 nM, 19.47 nM and 25.20 nM, respectively) (**Fig. S10-18**), likely due to increased electrostatic repulsion mediated by the inserted negatively charged residues. We next evaluated the *in vitro* cytotoxicity of the bone-targeting antibodies against HER2-positive and negative cell lines. Consistent with the flow cytometry data, Tras-CT, Tras-CH1/CT, and Tras-LC/CH1/CT antibodies kill SK-BR-3 cells with an efficiency similar to that of unmodified Tras (EC_50_ values of 5.97 ± 5.64 nM, 13.07 ± 12.09 nM, and 21.10 ± 20.25 nM, respectively) (**Fig. 2G)**. In contrast, none of the antibodies exhibit cytotoxicity against HER2-negative MDA-MB-468 cells under the same experimental conditions (**Fig. 2G**). These results indicate that introduction of the bone-homing sequence into antibodies can significantly enhance their bone affinity while preserving their anti-tumor activities.

### *In Vivo* Distribution of Bone-Targeting Antibodies

The effects of bone-homing peptides on Tras antibody distribution were further investigated in a mouse xenograft model. Using para-tibial injection, 2 × 10^5^ HER2-expressing MDA-MB-361 breast cancer cells labeled with firefly luciferase and red fluorescent protein were first introduced into the right leg of nude mice, followed by administration of sulfo-Cy7.5 labeled Tras, Tras-CT, Tras-CH1/CT, or Tras-LC/CH1/CT via retro-orbital injection. 72 h and 120 h after antibody infusion, the major organs were collected and imaged for antibody distribution. The intensity of the interosseous fluorescence signal was stronger in the Tras-CT, Tras-CH1/CT, and Tras-LC/CH1/CT injected animals than in Tras injected mice (**Fig. 2H, S20**, and **S21)**. Furthermore, larger quantities of bone-targeting antibodies were present in tumor-bearing right leg bones compared to left healthy bone, likely due to *L*-Asp_6_ peptide-mediated targeting to the bone resorption niche. Overall, these results indicate that introduction of the *L*-Asp_6_ sequence can significantly increase the concentration of therapeutic antibody in bone tumor sites. This effect has the potential to enhance antitumor activity at the tumor site, while at the same time decreasing systemic toxicity.

### *In Vivo* Therapeutic Activity of Bone-Targeting Antibodies against Bone Micrometastases

To determine whether bone-targeting antibodies can serve as novel therapeutic entities for the treatment of breast cancer metastasis to bone, we performed *in vivo* antitumor experiments in nude mice bearing MDA-MB-361 tumors. We inoculated 2 × 10^5^ MDA-MB-361 breast cancer cells labeled with firefly luciferase and red fluorescent protein into the right leg of nude mice via para-tibial injection. One week after injection, wild type Tras and bone-targeting Tras antibodies were administered by retro-orbital injection. As shown in **Fig. 3A**, mice receiving 1mg/kg of unmodified Tras did not respond well to this treatment. Despite an initial inhibitory effect during the first two weeks of treatment, unmodified Tras failed to control long-term tumor growth, prolonging median mouse survival by only 9.7 days (**Fig. 3B** and **C**). In contrast, when the tumor-bearing mice were treated with bone-targeting antibodies, tumor growth was significantly inhibited. Tras-CT-, Tras-CH1/CT-, and Tras-LC/CH1/CT-treated groups exhibited pronounced delays in tumor growth of 27.5, 35.8 and 18.9 days, respectively (**Fig. 3B** and **C**). **Figs. 3A, 3B, 3C, S22, S23**, and **Table S3** show the bioluminescent (BLI) signals for each treatment groups from day 1 to day 80. In the PBS-treated control group, there was a progressive increase in the BLI signal over time. The BLI signals from day 1 to 80 demonstrate that bone-targeting Tras antibody-treated groups experienced significant delays in tumor growth compared to the Tras-treated group (**Fig. 3B** and **C**). Mice treated with Tras-CH1/CT exhibited the smallest increases in tumor size (Tras-CH1/CT vs Tras: 9.3 ± 4.2 vs 1562.7 ± 801.6, *p* < 0.0001). Furthermore, the Tras-CH1/CT-treated mice experienced a 62.5% rate of survival, a significant improvement over that seen in Tras-treated mice (**Fig. 3D**). Thus, treatment with Tras-CH1/CT appears to result in more effective inhibition of micrometastasis progression than that seen in Tras-treated mice. Treatment with bone-targeting antibodies was well tolerated, with no overt signs of toxicity observed in any of the treatment groups. For example, we observed no differences in body weights across the various treatment groups (**Fig. 3E**).

**Figure 3.**
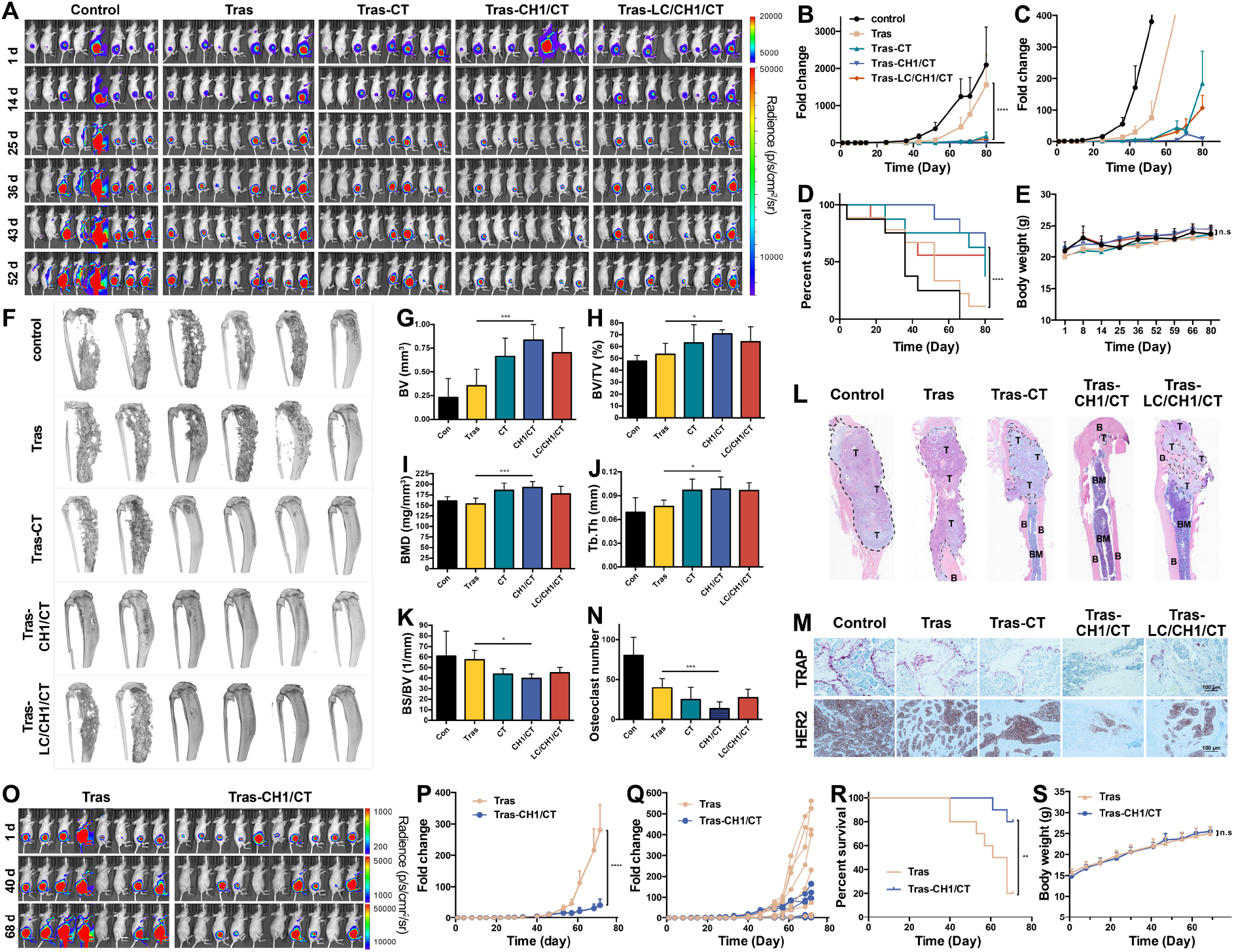
Bone-targeting antibodies inhibit breast cancer bone metastases. **(A)** MDA-MB-361 cells were para-tibia injected into the right hind limb of nude mice, followed by treatment with PBS, Tras (1 mg/kg retro-orbital venous sinus in sterile PBS twice a week), Tras-CT, Tras-CH1/CT, and Tras-LC/CH1/CT (same as Tras). Tumor burden was monitored by weekly bioluminescence imaging. **(B-C)** Fold-change in mean luminescent intensity of MDA-MB-361 tumors in mice treated as described in (A). *p* values are based on two-way ANOVA test. **(D)** Kaplan-Meier plot of the time-to-euthanasia of mice treated as described in (A). For each individual mouse, the BLI signal in the whole body reached 10^7^ photons sec^-1^ was considered as the endpoint. **(E)** Body weight change of tumor-bearing mice over time. **(F)** MicroCT scanning in the supine position for groups treated with PBS, Tras, Tras-CT, Tras-CH1/CT, and Tras-LC/CH1/CT. **(G)** Quantitative analysis of bone volume (BV). **(H)** Quantitative analysis of bone volume/tissue volume ratio (BV/TV). **(I)** Quantitative analysis of trabecular bone mineral density (BMD). **(J)** Quantitative analysis of trabecular thickness (Tb.Th). **(K)** Quantitative analysis of bone surface/bone volume ratio (BS/BV). **(L)** Representative longitudinal, midsagittal hematoxylin and eosin (H&E)-stained sections of tibia/femur from each group. T: tumor; B: bone; BM: bone marrow. **(M)** Representative images of HER2 and TRAP staining of bone sections from each group. **(N)** Osteoclast number per image calculated at the tumor-bone interface in each group (pink cells in (K) were considered as osteoclast positive cells). **(O)** MDA-MB-361 cells were para-tibia injected into the right hind limb of nude mice, followed by treatment with Tras (10 mg/kg retro-orbital venous sinus in sterile PBS every two weeks) and Tras-CH1/CT (same as Tras). Tumor burden was monitored by weekly bioluminescence imaging. **(P)** Fold-change in mean luminescent intensity of MDA-MB-361 tumors in mice treated as described in (O). **(Q)** Fold-change in individual luminescent intensity of MDA-MB-361 tumors in mice treated as described in (O). **(R)** Kaplan-Meier plot of the time-to-euthanasia of mice treated as described in (O). For each individual mouse, the BLI signal in the whole body reached 10^8^ photons sec^-1^ was considered as the endpoint. **(S)** Body weight change of tumor-bearing mice in (O) over time. \*\*\*\**P* < 0.0001, \*\*\**P* < 0.001, \*\**P* < 0.01, \**P* < 0.05, and n.s. *P* > 0.05.

At the end of the experiment (day 81), tibiae (from tumor-bearing legs) were harvested and scanned by micro-computed tomography (mico-CT). The micro CT analysis revealed extensive osteolytic bone destructions in both the PBS- and Tras-treated groups, while bone loss was significantly reduced in bone-targeting antibody-treated groups (**Fig. 3F, S24** and **S25**). Compared to mice in the PBS- and Tras-treated groups, Tras-CH1/CT-treated mice exhibited greater bone volume (BV, **Fig. 3G**), greater bone volume/tissue volume ratio (BV/TV, **Fig. 3H**), greater bone mineral density (BMD, **Fig. 3I**), and thicker trabecular bone (Tb.Th, **Fig. 3J**), but smaller bone surface/bone volume ratio (BS/BV, **Fig. 3K** and **Table S4**). These parameters are all indicative of significant retardation of micrometastasis-induced osteolysis by antibodies modified with bone-homing peptides. Histological analysis further reveals the invasion of tumor cells into the bone matrix and into the adjacent tissue in the Tras-treated group (**Fig. 3L**). Histology also confirms the reduction of intratibia tumor burden in these mice that was indicated by the BLI results. Bone sections from Tras-CH1/CT treated mice reveal the reduced tumor growth and relatively normal bone morphology in these mice, consistent with significant inhibition of tumor invasion and bone destruction (**Fig. 3L**). Bone samples from the various treatment groups were also analyzed for bone resorbing TRAP (tartrateresistant acid phosphatase) positive multinucleated osteoclasts (shown as pink cells) and HER-expressing cancer cells (**Fig. 3M, 3N, S26-28**). Compared to Tras treatment, Tras-CH1/CT treatment significantly reduces the numbers of both osteoclasts and HER2-positive cells in bone niches, once again consistent with the ability of bone targeting antibodies to inhibit micrometastasis progression (**Fig. 3N**). Given that tumor-induced hypercalcemia and TRACP 5b protein are the indicators of osteolytic bone destruction, the effects of bone-targeting antibody-treated were evaluated. The results suggested that Tras-CH1/CT-treated groups has relative better therapeutic effects (**Fig. S29**). Next, we evaluated the benefits of bone-targeting antibodies for bone metastasis at a high dose. Nude mice with MDA-MB-361 tumors were treated with Tras or Tras-CH1/CT at 10 mg/kg every two weeks. Notably, Tras-CH1/CT treatment results in statistically significant growth inhibition, compared to that seen in mice treated with 10 mg/kg of Tras (**Fig. 3O, 3P, 3Q, S30 and S31**). As shown in Fig. 3R, mice injected with Tras-CH1/CT at 10 mg/kg had significantly longer overall survival than did mice injected with the same amount of Tras (**Fig. 3R**). Furthermore, no weight loss was observed with a high dose of bone-targeting antibodies (**Fig. 3S**).

To evaluate the enhanced therapeutic efficacy of bone-targeting antibodies in a second tumor model, we next treated nude mice implanted with MCF-7 breast cancer cells. Animals were then treated with Tras or Tras-CH1/CT at 1 mg/kg twice a week. Consistent with the results obtained in the MDA-MB-361 model, significant reductions in tumor BLI signals and prolonged survival of mice were observed in the Tras-CH1/CT-treated group, compared to values found in the Tras-treated group (**Figs. 4A, 4B, 4C, 4D** and **S32**). Tras-CH1/CT treatment did not induce any weight loss in animals (**Fig. 4E**).

**Figure 4.**
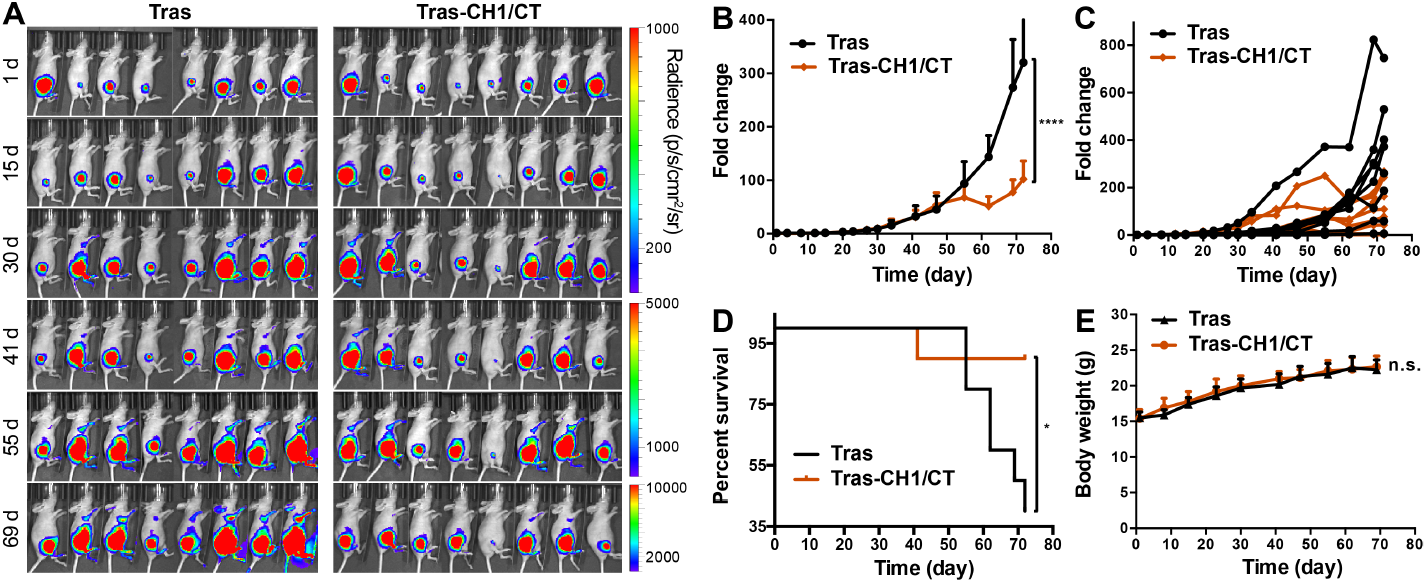
**(A)** MCF-7 cells were para-tibia injected into the right hind limb of nude mice, followed by treatment with Tras (1 mg/kg retro-orbital venous sinus in sterile PBS twice a week) and Tras-CH1/CT (same as Tras). Tumor burden was monitored by weekly bioluminescence imaging. **(B)** Fold-change in mean luminescent intensity of MCF-7 tumors in mice treated as described in (A). *p* values are based on two-way ANOVA test. **(C)** Fold-change in individual luminescent intensity of MCF-7 tumors in mice treated as described in (A). **(D)** Kaplan-Meier plot of the time-to-euthanasia of mice treated as described in (A). For each individual mouse, the BLI signal in the whole body reached 5×10^7^ photons sec^-1^ was considered as the endpoint. **(E)** Body weight change of tumor-bearing mice in (A) over time. \*\*\*\**P* < 0.0001, \**P* < 0.05, and n.s. *P* > 0.05.

### Bone-Targeting Antibodies Inhibit Secondary Metastases from Bone Lesions

In more than two-thirds of patients, breast cancer metastases are not restricted to the skeleton, but subsequently also occur in other organs.^28,29,30,31^ Recent genomic analyses suggest that these metastases, the major cause of morbidity and mortality, are not derived from primary tumors, but are seeded from other metastatic sites. Taking advantage of a recently developed approach that selectively delivers cancer cells to hind limb bones, we and others have observed frequent “metastasis-to-metastasis” seeding from established bone lesions to multiple other organs (**Fig. 5A**).^32–35^ Hence, it is imperative for us to evaluate whether bone-targeting antibodies can inhibit these secondary metastases derived from bone lesions. Using the para-tibiae injection method, we can obtain highly bone-specific tibiae-bearing tumors at early stages of development. As the bone lesion progresses, metastases marked by bioluminescence signals began to appear in many other organs, including other bones, lungs, heart, liver, spleen, kidney, and brain. Using this model, 2 × 10^5^ luciferase-labeled MDA-MB-361 cells were introduced into the right hind limbs of nude mice via para-tibial injection, followed by treatment with unmodified Tras and bone-targeting Tras antibodies. Mice were euthanized 81 days after initiation of treatment, and organs were harvested for *ex vivo* bioluminescence imaging. Seven organs including the right hindlimb bone and six soft tissue organs were isolated for assessment of metastasis. As shown in **Fig. 5B, 5C** and **S33**, treatment with Tras-CH1/CT significantly reduces the frequency of secondary metastasis to the contralateral hind limb (left hindlimb bone), heart, and liver. The *ex vivo* BLI intensities of secondary metastases from primary bone lesions are reduced in experimental groups treated with bone-targeting antibodies, especially in the Tras-CH1/CT-treated group. Compared to treatment with unmodified Tras, the metastatic signals from the lung and liver were significantly decreased after the Tras-CH1/CT treatment (**Fig. 5C and S33**). Taken together, these data reveal the enhanced therapeutic efficacy of bone-targeting antibodies against both initial bone metastasis and secondary metastasis from the bone to other distant organs.

**Figure 5.**
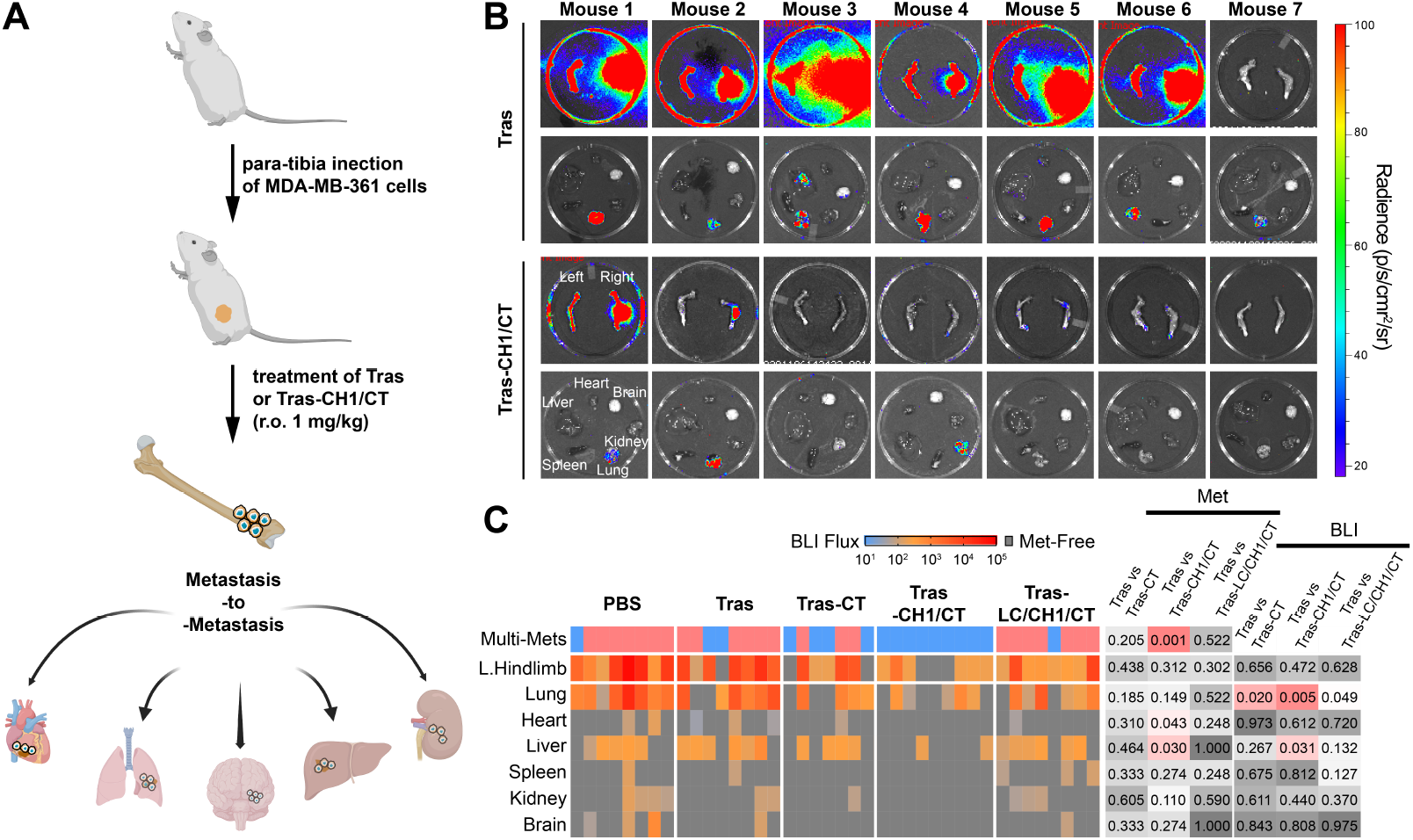
**(A)** Bone lesions more readily give rise to secondary metastases to multiple organs. **(B)** Secondary metastases observed in various organs in mice treated with Tras or Tras-CH1/CT. **(C)** Heat map of *ex vivo* BLI intensity and status of metastatic involvement in tissues from mice treated with PBS, Tras, Tras-CT, Tras-CH1/CT, and Tras-LC/CH1/CT. Each column represents an individual animal, and each row represents a type of tissue. The presence of the metastasis was defined as the presence of BLI signal above 18 counts/pixel under 120 seconds exposure time. Multi-site metastases were defined as the metastatic involvement of at least three tissues. p values were determined by Fisher’s exact test on the frequency of metastatic involvement while by Mann-Whitney test of on the metastatic burden.

### Modification of Antibody-Drug Conjugates with the Bone-Homing Peptide Exhibit Enhanced Therapeutic Efficacy in vivo

Antibody-drug conjugates (ADCs) that combine the antibody’s tumor specificity with the high toxicity of chemotherapy drugs, are emerging as an important class of anticancer drugs for breast cancer patients, especially ones with advanced breast cancer.^36,37^ Following the first FDA approval of trastuzumab emtansine (T-DM1) for HER2-positive breast cancer, trastuzumab deruxtecan has been recently approved for the treatment of adults with unresectable or metastatic HER2-positive breast cancers.^38,39,40^ It is possible that adding the bone-specificity to ADCs can improve their efficacy against breast cancer bone metastasis. To test this possibility, we first used pClick conjugation technology to site-specifically couple the monomethyl auristatin E (MMAE) to both wild type antibody Tras and the bone-targeting antibody Tras-CH1/CT with the best anti-cancer activity (**Fig. 6A**).^41,42^ The successful preparation of the conjugates was demonstrated by SDS–polyacrylamide gel electrophoresis (PAGE) and electrospray ionization mass spectrometry (ESI-MS) (**Fig. 6B, S34** and **S35**). To ensure that conjugation of toxin did not alter the antigen targeting ability and specificity, the in vitro binding assays were performed using HER2-positive and -negative cells (**Fig. S36**). Tras-CH1/CT-MMAE showed a high binding affinity to HER2-positive SK-BR-3 cells, but not HER2-negative MDA-MB-468 cells. Next, we evaluated the in vitro cytotoxicity of these ADCs in SK-BR-3 and MDA-MB-468 breast cancer cell lines (**Fig. 6C** and **6D**). Both Tras-MMAE and Tras-CH1/CT-MMAE exhibited high potency only in the SK-BR-3 (EC50: 0.18 ± 0.82 nM and 0.49 ± 0.30 nM, respectively), with no significant toxicity was observed in the MDA-MB-468 cells. To test the bone targeting ability of Tras-CH1/CT-MMAE *in vitro*, we incubated either Tras-CH1/CT-MMAE or Tras-MMAE with nondecalcified bone sections from C57BL/6 mice. As expected, only the signal of Tras-CH1/CT-MMAE correlated well with the XO signal, confirming its bone-targeting ability (**Fig. 6E**).

**Figure 6.**
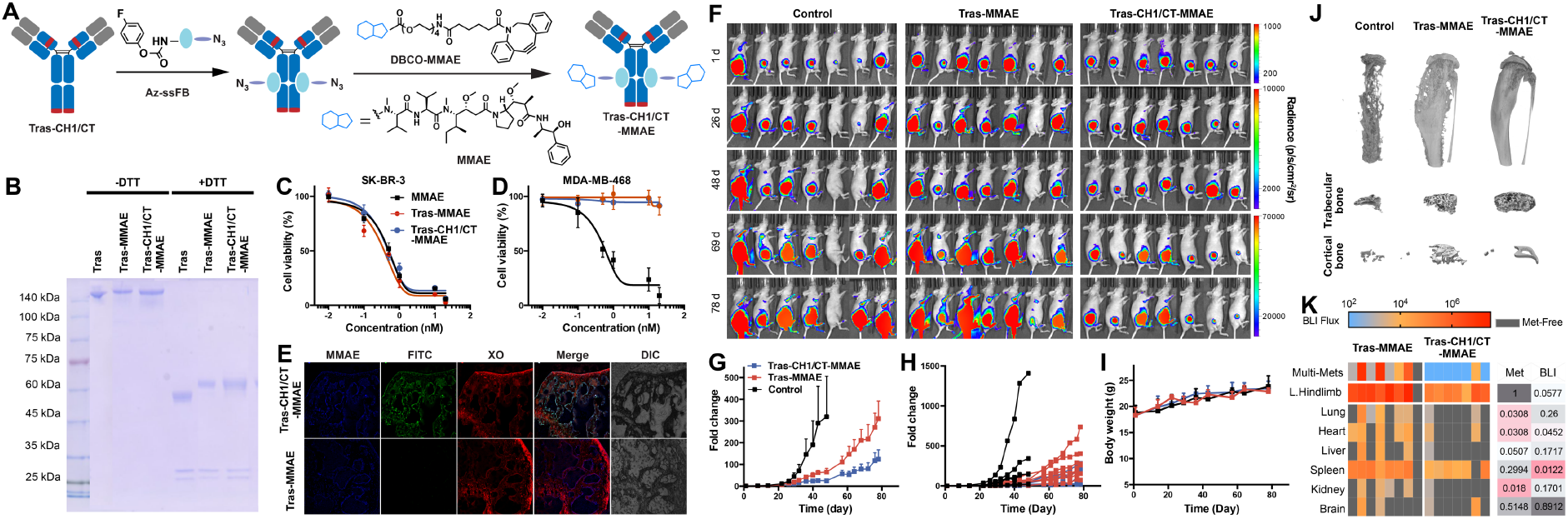
**(A)** Preparation of bone-targeting antibody-drug conjugate. Tras antibody was first modified with the bone-homing peptide at the heavy chain (CH1) and c-terminus (CT), followed by the modification of MMAE using pClick antibody conjugation technology. **(B)** SDS-PAGE analysis of Tras-MMAE and Tras-CH1/CT-MMAE under non-reducing (left) and reducing (right) conditions. **(C-D)** In vitro cytotoxicity of MMAE, Tras-MMAE, and Tras-CH1/CT-MMAE, against SK-BR-3 and MDA-MB-468 cancer cells. **(E)** Differential bone targeting ability of Tras-MMAE and Tras-CH1/CT-MMAE. Nondecalcified bone sections from C57/BL6 mice were incubated with 50 µg/mL Tras-MMAE and Tras-CH1/CT-MMAE overnight, followed by staining with fluorescein isothiocyanate (FITC)-labeled anti-human IgG and 4 µg/mL xylenol orange (XO, known to label bone). **(F)** MDA-MB-361 cells were para-tibia injected into the right hind limb of nude mice, followed by treatment with Tras-MMAE (0.5 mg/kg retro-orbital venous sinus in sterile PBS every week) and Tras-CH1/CT-MMAE (same as Tras). Tumor burden was monitored by weekly bioluminescence imaging. **(G)** Fold-change in mean luminescent intensity of MDA-MB-361 tumors in mice treated as described in (F). **(H)** Fold-change in individual luminescent intensity of MDA-MB-361 tumors in mice treated as described in (F). **(I)** Body weight change of tumor-bearing mice in (F) over time. **(J)** MicroCT scanning in the supine position for groups treated with Tras-MMAE and Tras-CH1/CT-MMAE after tumor implantation. **(K)** Heat map of *ex vivo* BLI intensity and status of metastatic involvement in tissues from mice treated with Tras-MMAE and Tras-CH1/CT-MMAE. Each column represents an individual animal, and each row represents a type of tissue. The presence of the metastasis was defined as the presence of BLI signal above 18 counts/pixel under 120 seconds exposure time. Multi-site metastases were defined as the metastatic involvement of at least three tissues. *p* values were determined by Fisher’s exact test on the frequency of metastatic involvement while by Mann-Whitney test of on the metastatic burden. ^****^*P* < 0.0001, ^***^*P* < 0.001, ^**^*P* < 0.01, ^*^*P* < 0.05, and n.s. *P* > 0.05.

Next, we examined the therapeutic efficacy of bone-targeting ADCs in the xenograft model of bone metastasis. To our delight, weekly administration of the Tras-CH1/CT-MMAE led to a significant inhibition effect, compare to unmodified Tras-MMAE (**Fig. 6F-H, S37 and S38**). Furthermore, the bone-targeting ADC showed no appearance toxicity, with a continuous increase of body weight across the different groups during treatment (**Fig. 6I**). The micro-CT analysis revealed extensive osteolytic bone destructions in both the PBS- and Tras-MMAE-treated group, while bone loss was significantly reduced in Tras-CH1/CT-MMAE-treated group (**Fig. 6J** and **S39**). Compared to mice in the PBS- and Tras-MMAE-treated groups, Tras-CH1/CT-MMAE-treated mice exhibited higher bone volume/tissue volume ratio (BV/TV, **Fig. S40A**) and thicker trabecular bone (Tb.Th, **Fig. S40B**). Histology also confirms the reduction of intratibia tumor burden in Tras-CH1/CT-MMAE mice that was indicated by the BLI results (**Fig. S41**). Furthermore, we observed that bone-targeting ADC could better inhibit these secondary metastases derived from bone lesions. Much higher levels of lung, heart, and kidney metastasis were detected in Tras-MMAE group, compare to Tras-CH1/CT-MMAE treatment groups (**Fig. 6K** and **S42**). Taken together, these results indicate that bone-specific delivery of ADCs to bone micrometastasis sites will also benefit breast cancer patients with bone metastases.

## DISCUSSION

Antibody therapy has evolved to focus on finding new biomarkers and on functionalizing antibodies with new payloads. We now demonstrate that adding tissue homing capability to therapeutic antibodies can improve their tumor-specific distribution, thereby enhancing therapeutic efficacy. Failure to achieve an efficacious dose in the tissue of interest following intravenous injection is a major shortcoming of many antibody-based therapeutics. Incomplete access of therapeutics to all cells in tumor tissues can lead to treatment failure and to the development of acquired drug resistance. As an example, therapeutic antibodies that exhibit excellent efficacy in the treatment of primary tumors often yield suboptimal responses against bone or brain metastases that offer limited access to macromolecules. Furthermore, antibody-based therapeutics can be associated with unacceptable ‘on-target’ toxicity for cases in which target antigens are also present in healthy tissues.^43^ For example, the use of trastuzumab for breast cancers that overexpress HER2 is associated with rare but fatal lung toxicity, referred to as interstitial lung disease. Therefore, adding tissue specificity to improve selective delivery of antibodies to tumors provides a promising avenue for advancing new anticancer therapies toward clinical translation.

Antibody-based therapy has been approved for use in adult patients with metastatic breast cancers, but the performance of this regimen in patients with bone metastases has been disappointing. Antibody-based trastuzumab (Herceptin) and pertuzumab (Perjeta) therapies are established standards of care for HER2+ adjuvant and metastatic breast cancer. Although many HER2+ bone metastatic breast cancer patients benefit from these treatments, few experience prolonged remission.^16,44–46^ Of patients with HER2-positive bone metastases, only 17% achieved a complete response and none achieved a durable complete response. By comparison, 40% and 30% of patients with liver metastases achieved complete responses and durable complete responses, respectively. As a further example, the immune checkpoint inhibitors atezolizumab (Tecentriq) and pembrolizumab (Keytruda) are anti-PD-L1 antibodies that are approved by the Food and Drug Administration for treatment of metastatic triple-negative breast cancer. However, a recent phase III clinical trial revealed that these drugs do not benefit patients with bone metastases. Among breast cancer patients with bone metastases, no significant difference was observed between atezolizumab and placebo groups for the risk of progression or death (median progression-free survival, 5.7 months vs. 5.2 months, stratified hazard ratio for progression or death, 1.02; 95% CI, 0.79-1.31).^47^ Thus, strategies for improving the outcomes of breast cancer patients with bone metastases are highly desired.

In this study, we demonstrate that adding bone-homing peptides to therapeutic antibodies leads to increased antibody concentrations in the bone metastatic niche, relative to other tissues. This approach yields a targeted therapy against bone metastases as well as against secondary multi-organ metastatic seeding from bone lesions. Using xenograft models of bone metastasis, we find that unmodified trastuzumab has poor bone tissue penetration and distribution, thus reducing access of the antibody to its target and limiting its efficacy against cancer cells in the bone microenvironment. Compared with the unmodified antibody, trastuzumab modified with bone-homing peptide sequences (*L*-Asp_6_) exhibit enhanced targeting to sites of bone metastasis. This results in improved activity against breast cancer metastasis to the bone and against metastatic seeding from bone lesions to other organs. Most importantly, we demonstrate that the modified antibody and antibody-drug conjugate with moderate bone-binding capability has optimal efficacy *in vivo*. In contrast, an antibody with higher bone-binding capability has suboptimal activity against bone metastases. This may be due to slow release of the latter entity from the bone matrix or to increased electrostatic repulsion resulting from the increased number of peptides with negative charges. The addition of bone specificity to antibody therapy enables the specific delivery of these agents to the bone. This study establishes a new strategy for transitioning antibody-based therapies from antigen-specific to both antigen and tissue-specific. The strategy not only results in enhanced therapeutic efficacy, but may also reduce adverse side effects associated with systemic distribution of the drug, thus providing a promising new avenue for advancing antibody therapy toward clinical translation.

## MATERIALS AND METHODS

### Materials

Unless otherwise noted, the chemicals and solvents used were of analytical grade and were used as received from commercial sources. LB agar was ordered from Fisher. Oligonucleotide primers were purchased from Eurofins Genomics (Supplementary **Table S1**). Plasmid DNA preparation was carried out with the GenCatchTM Plus Plasmid DNA Miniprep Kit and GenCatchTM Advanced Gel Extraction Kit. PNGase F was purchased from New England Biolabs to remove the glycans before ESI-MS analysis. NuPAGE 4-12% Bis-Tris Gel was purchased from Invitrogen. SDS-PAGE Sample Loading Buffer [6x] was purchased from BIOSCIENCES. PM2500 ExcelBand 3-color Regular Range Protein Marker was purchased from SMOBIO. ExpiCHO Expression Medium, ExpiFectamine CHO Transfection Kit, OptiPRO SFM Complexation Medium were purchased from Thermofisher. Hoechst 33342 (Cat No: H1399) were purchased from Life TechnologiesTM. 3,3-Dioctadecyloxacarbocyanine perchlorate (DiIC18, Cat No: M1197) was purchased from Marker Gene Technologies, Inc.

### Cell lines

MDA-MB-361, BT474, SK-BR-3, MCF-7, and MDA-MB-468 cell lines were cultured according to ATCC instructions. Firefly luciferase and RFP labeled MDA-MB-361 cell line was generated as previously described.^48^

### SDS-PAGE analysis

The intact and reduced antibody samples were analyzed using invitrogen NuPAGE 4-12% Bis-Tris gel. 20 μL of 0.2 mg/mL antibody samples were mixed with 4 μL sample loading dye before loading into the gel. The gel was running under 160 V in MES buffer for 35 min and stained using commassie blue buffer. The gel images were taken by Amersham Imager 600 and analyzed by ImageQuanTL software.

### ESI-MS analysis

The antibodies were analyzed using a single quadrupole mass spectrometer (Agilent: G7129A) coupled with 1260 infinity II Quaternary Pump (Agilent: G7111B). (Column: Pursuit 5 Diphenyl 150 × 2.0 mm) The elution conditions for antibodies were as follows: mobile phase A= 0.1% formic acid water; mobile phase B= 0.1% formic acid in acetonitrile; gradient 0–0.1 min, 10–15% B; 0.1–8 min, 15–50% B; 8–8.1 min, 50–10% B; flow rate= 0.5 mL/min. The absorbance was measured at 280 nm. Automatic data processing was performed with MassHunter BioConfirm software (Agilent) to analyze the intact and reduced MS spectra.

### HA binding assay

Briefly, 1 mg of Tras or bone-targeting antibodies were diluted in 0.5 mL PBS (pH 7.4) in an Eppendorf tube. HA (20 equiv, 20 mg) was suspension in 0.5 mL PBS. Then, the antibodies and HA were mixed with vortex, and the resulting suspension was shaken at 220 rpm at 37 ºC. Samples without HA were used as controls. After 0.25, 0.5, 1, 2, 3, 6 and 8 hours, the suspension was centrifuged (3000 rpm, 3 min) and the absorbance of the supernatant at 280 nm was measured by Nanodrop. The percent binding to HA was calculated as follow, where OD represents optical density:

[(OD_without HA_ – OD_with HA_)/(OD_without HA_)] × 100%.

### *In vitro* cytotoxicity of bone-targeting antibodies and bone-targeting ADC

SK-BR-3 and MDA-MB-468 cells at 2 × 10^3^ cells/well into 96-well plates. After 24 h incubation, cells were treated with different concentrations of Tras, Tras-CT, Tras-CH1/CT and Tras-LC/CH1/CT, and then incubated for 4 d (Tras, Tras-CT, Tras-CH1/CT and Tras-LC/CH1/CT) or 3 d (MMAE, Tras-MMAE and Tras-CH1/CT-MMAE). 20 *μ*L of 3-(4,5-dimethylthiazol-2-yl)-2,5-diphenyltetrazolium bromide (MTT) solution (5 mg/mL) was then added to each well and incubated for another 4 h. Medium was aspirated and 150 *μ*L DMSO was added to each well. The absorbance at 565 nm was measured by microplate reader (Infinite M Plex by Tecan) to quantify living cells.

### Flow cytometry

Cancer cells (3 × 10^5^) were incubated with 30 μg/mL Tras and bone-targeting antibodies for 30 min at 4°C. After washing away unbound antibodies, bound antibodies were detected using Fluorescein (FITC) AffiniPure Goat Anti-Human IgG (H+L) (code: 109-095-003, Jackson Immunology) for 30 min at 4°C. Fluorescence intensity was determined using a BD FACSVerse (BD Biosciences).

### Confocal imaging

Cancer cells were grown to about 80% confluency in eight-well confocal imaging chamber plates. The cells were incubated with 30 nM FITC labeled antibodies for 30 min and then fixed by 4% paraformaldehyde for 15 min. The cells were washed three times with PBS (pH 7.4) and then incubated Hoechst 33342 (catalog number H1399, Life Technologies) for 5 min. The cells were then washed three times with PBS (pH 7.4) and used for confocal imaging. Confocal fluorescence images of cells were obtained using a Nikon A1R-si Laser Scanning Confocal Microscope (Japan).

### Construction of Tras-CH1/CT-MMAE

ssFB-N_3_ was prepared as previously reported.^**49**^ In briefly, 20 equiv of ssFB-N_3_ peptide (0.4 mM) was mixed with mAb (0.5 mg/mL in PBS) at 37 °C for two days. The mAb-FB-N_3_ conjugate was purified via a PD-10 desalt column. 40 equiv of DBCO-(PEG)4-MMAE was added at RT overnight for strain-promoted alkyne-azide cycloaddition (SPAAC) reaction. Finally, Tras-MMAE or Tras-CH1/CT-MMAE were purified via a PD-10 desalt column and characterized by ESI-MS.

### Determination of *K*_*d*_ values

The functional affinity of bone-targeting antibodies for HER2 was determined as reported.^50^ Briefly, a total of 2 × 10^5^ SK-BR-3 or MDA-MB-468 cells were incubated with graded concentrations of Tras, Tras-LC, Tras-CT, Tras-CH1, Tras-LC/CT, Tras-CH1/CT, Tras-LC/CH1, and Tras-LC/CH1/CT for 4 hours on ice. Then, the bound antibody was detected by Fluorescein (FITC) AffiniPure Goat Anti-Human IgG (H+L) (Jackson Immunology). Cells were analyzed for fluorescence intensity after propidium iodide (Molecular Probes, Eugene, OR) staining. The linear portion of the saturation curve was used to calculate the dissociation constant, KD, using the Lineweaver-Burk method of plotting the inverse of the median fluorescence as a function of the inverse of the antibody concentration. The KD was determined as follows: 1/F=1/Fmax+(*K*_*d*_/Fmax)(1/[Ab]), where F corresponds to the background subtracted median fluorescence and Fmax was calculated from the plot.

### Binding to bone cryosections

Nondecalcified long bone sections from C57BL/6 mice were incubated with 50 μg/mL Tras, Tras-CT, Tras-CH1/CT, Tras-LC/CH1/CT, Tras-MMAE or Tras-CH1/CT-MMAE, conjugated overnight at 4 °C, followed by staining with fluorescein isothiocyanate (FITC)-labeled anti-human IgG for 60 min at room temperature. After washing 3 times with PBS, specimens were incubated for 30 min at 37 °C with Xylenol Orange (XO) (stock: 2 mg/ml, dilute 1:500, dilute buffer: PBS pH 6.5). After three washes with PBS, specimens were stained with Hoechst 33342 (stock 10mg/ml, dilute 1:2000) for 10 min. Slides were then washed with PBS, air dried, and sealed with Prolong™ gold anti-fade mountant (from ThermoFisher).

### *In vivo* evaluation of Tras, Tras-CT, Tras-CH1/CT, and Tras-LC/CH1/CT

To establish bone metastasis, firefly luciferase and RFP labeled MDA-MB-361 cells (2 × 10^5^) or MCF-7 cells (2 × 10^5^) were inoculated into tibia of 3-4 weeks old female athymic nude mice using para-tibial injection method. 7 days after surgery, mice were ranked/random divided to obtain similar tumor burden in each group. PBS and antibodies were injected via retro-orbital injection. Animals were imaged once a week using IVIS Lumina II (Advanced Molecular Vision), following the recommended procedures and manufacturer’s settings. All of the mice were euthanized after blood was collected on day 81, and all the organs (tumor-bearing tibia, heart, liver, spleen, lung, brain and kidney) were collected for further tests.

### *Ex vivo* metastasis-to-metastasis analysis

At the end point, live animals were given D-Luciferin and immediately dissected. The tissues were examined by *ex vivo* BLI imaging. The whole process organ collection procedure and *ex vivo* imaging process should be finished within 15 minutes for each mouse.

### Bone histology and immunohistochemistry

At the end of experiments, mice were euthanized, and tibiae were harvested, fixed and then decalcified in 12% EDTA for 10 days. The tibiae were embedded in paraffin, and sectioned. Tumor burden was evaluated on hematoxylin and eosin (H&E) sections. Osteoclasts within the tumor and on bone-tumor-interface were counted after staining with Acid Phosphatase, Leukocyte (TRAP) Kit (Sigma). Immunohistochemistry analysis was performed on decalcified paraffin-embedded tissue sections using the HRP/DAB ABC IHC KIT (abcam) following the manufacture’s protocol.

### Radiographic analysis

Tibiae were dissected, fixed and scanned by microcomputed tomography (micro-CT, Skyscan 1272, Aartselaar, Belgium) at a resolution of 16.16 μm/pixel. Raw images were reconstructed in NReconn and analyzed in CTan (SkyScan, Aartselaar, Belgium) using a region of interest (ROI). Bone parameters analyzed included trabecular thickness (Tb.Th), bone volume fraction (BV/TV), bone mineral density (BMD), and bone surface/bone volume ratio (BS/BV).

### Biodistribution

Female athymic nude mice were injected para-tibia with MDA-MB-361 cells (2 × 10^5^ cells/animal). After 80 days, Cy7.5 labeled Tras and bone-targeting antibodies were administrated by retro-orbital injection. After 72 h and 120 h injection, the mice were imaged using IVIS. 72 h and 120 h injection, the mice were killed, and major organs including heart, liver, spleen, kidney, lung, and bone tumor tissue were removed. The fluorescence intensity in organs and tumor bearing tibiae were observed using IVIS.

For the wild type mice, Cy7.5 labeled Tras, Tras-CT and Tras-LC/CH1/CT were administered via retro-orbital injection to the C57BL/6 mice. After 48 hours, organs were dissected and imaged using IVIS.

### Quantification of TRAP and calcium levels in serum

At the end of experiments, blood was collected by cardiac puncture, and centrifuged for 15 min at 3,000 rpm to obtain the serum. The concentration of osteoclast-derived TRACP 5b was measured by using a Mouse ACP5/TRAP ELISA Kit (catalog number IT5180, GBiosciences). Serum calcium levels were determined colorimetrically using a calcium detection kit (catalog number DICA-500, Bioassays).

### Statistical methods

Data are presented as means plus or minus SEM and statistically analyzed using GraphPad Prism software version 6 (GraphPad software, San Diego, CA). Two-way ANOVA followed by Sidak’s multiple comparisons was used for all data collected over a time course. One-way ANOVA followed by Tukey’s multiple comparisons was used for Micro-CT data. Unpaired Student’s *t*-test was used for multi-organ metastasis data. *P* < 0.05 was considered to represent statistical significance.

## Acknowledgments

We thank Dr. Xiao and Zhang Laboratory members for insightful comments. This work was supported by the Cancer Prevention Research Institute of Texas (CPRIT RR170014 to H.X.), NIH (R35-GM133706 to H.X., R21-CA255894 to H.X., CA183878 and CA221946 to X.H.-F.Z.), the Robert A. Welch Foundation (C-1970 to H.X.), US Department of Defense (BC201371 to H.X. and X.H.-F.Z., DAMD W81XWH-16-1-0073, Era of Hope Scholarship to X.H.-F.Z.), the John S. Dunn Foundation Collaborative Research Award (to H.X.), and the Hamill Innovation Award (to H.X.). H.X. is a Cancer Prevention & Research Institute of Texas (CPRIT) scholar in cancer research.

## Author contributions

Z.T., C.Y., X.H.-F.Z. and H.X. developed the hypothesis, designed experiments, analyzed the data, and wrote the manuscript. Z.T., C.Y., W.Z, K.W., R.G., Z.X. and L.W. performed experiments. W.Z., Y.C., X.H.-F.Z. and H.X. contributed to experimental design, generation of the reagents, and manuscript editing. X.H.-F.Z. and H.X. conceived and supervised the project.

